# Iron sensing modules participate in rhizospheric pH perception in rice

**DOI:** 10.64898/2026.10.23.761470

**Authors:** Ajit Pal Singh, Hamida Bano, Reshma Gopal, Ajay K Pandey

**Author notes:** **Author for correspondence:** Ajit Pal Singh, Department of Biotechnology, Sharda School of Bio-Science and Technology, Sharda University, Greater Noida, Uttar Pradesh-201310, India.

## Abstract

Rhizospheric pH has emerged as one of the most important environmental factors regulating plant fitness in almost every aspect of plant physiology and development. Interestingly, the information on how plant perceive rhizospheric pH is still not well understood. In this report we have elucidated the intricate mechanism where rhizospheric pH mediated changes in relative Fe availability are sensed by the plants to initiate complex physiological responses. Using root specific RNA-seq, ICPMS, hormone quantification and physiological validation in rice, the study revealed Fe availability to be a critical determinant of rhizospheric pH where the relative Fe toxicity towards acidic pH and Fe deficiency towards alkaline pH is deduced by the plant in the form of JA-auxin crosstalk regulating the physiological responses against rhizospheric pH.

## Introduction

The rhizosphere, defined as the narrow zone of soil in direct contact with plant roots, is a dynamic microenvironment where roots, soil particles and microorganisms interact to shape plant growth and nutrient acquisition. Rhizospheric pH (pH_Ext_) plays a pivotal role in directing a plethora of processes including nutrient availability, root growth and dynamicity of microbial community (**Gregory & Hinsinger, 1999; Fierer and Jackson, 2006; Rousk et al., 2010**). While most of the plant only survive over a very narrow range of pH_Ext_, they do so by maintaining a relatively stable pH range at the cytoplasmic as well as organelle level (**Arnon and Johnson, 1942; Tsai and Schmidt, 2021**). The importance of understanding plant responses to pH_Ext_ is underscored by its global agricultural relevance (**von Uexküll and Mutert, 1995**). For an estimate around 40% of the world’s arable land is characterized as acidic soil and around 1 billion hectare falls under alkaline pH category (**Zhang et al., 2023**). As environmental conditions such as flooding, drought and soil degradation continue to change soil pH, dynamics are expected to become even more variable, making it essential to understand how plants respond to these fluctuations. Moreover, improving crop performance under suboptimal pH conditions is a major challenge for sustainable agriculture, particularly in the context of nutrient use efficiency and stress tolerance.

Root growth is strongly influenced by pH-dependent cell wall dynamics, where acidic condition is known to promote cell elongation (consistent with the acid growth hypothesis), while alkaline conditions restrict cell elongation and require compensatory mechanisms such as cytoskeletal reorganization (**Qian et al., 2024; Singh et al., 2025**). Interestingly, despite the cell elongation on acidic pH and inhibition on alkaline pH, root growth inhibition remains a common response against changing rhizospheric pH towards both the extremes. Importantly, rhizospheric pH exerts its developmental influence largely through modulation of hormone signaling, particularly auxin and jasmonate (**Singh et al., 2025**). Interestingly, auxin transport is also known to be dependent on proton gradients across the plasma membrane and therefore changes in external pH can directly alter auxin distribution within root tissues (**Lager et al., 2010; Singh et al., 2025**). Auxin in turn has significant consequences for root system architecture, including lateral root elongation, as demonstrated in both rice and Arabidopsis systems (**Rahman et al., 2002**; **Wang et al., 2003**; **Nakamura et al., 2023**). At the cellular level, root hair development is tightly associated with localized oscillations in apoplastic pH and reactive oxygen species (ROS), which together regulate tip growth and expansion (**Bibikova et al., 1998**; **Monshausen et al., 2007**).

Beyond development, pH_Ext_ critically determines nutrient availability and uptake, making it a key factor in plant nutrition. The solubility and speciation of essential nutrients such as iron (Fe), and other micronutrients are highly pH-dependent (**Fageria and Zimmermann, 1998; Yang et al., 2026**). Early foundational work has demonstrated that nutrient uptake patterns can significantly alter the soil pH surrounding roots and *vice-versa* (**Ortas and Rowell, 2004**). More recent studies have reinforced that these pH mediated changes are not merely passive consequences of metabolism but are tightly regulated processes including plant physiology. For instance, rapid transcriptional responses to pH_Ext_ shifts have been observed, indicating that plants perceive and respond to rhizospheric pH (**Lager et al., 2010; Bailey et al., 2023**).

Despite its importance, the mechanisms of pH_Ext_ perception in plants remain poorly understood. One major challenge is the lack of clearly identified molecular mechanism of pH sensing. Unlike other environmental signals such as light or hormones, which have well-characterized receptors, the mechanism of pH perception in plants has not yet been identified (**Tsai and Schmidt, 2021**). Several emerging perspectives are shaping current understanding of pH perception in plants. One perspective proposes that pH acts as an integrative signal reflecting multiple aspects of the plant’s environment including nutrient status and stress conditions, simultaneously. Another perspective is the distributed sensing model, in which pH perception arises from the collective activity of multiple sensors and pathways, rather than a single receptor for pH_Ext_ perception. There is also increasing recognition of the importance of hormonal crosstalk, with pH influencing or being influenced by signaling networks involving auxin, jasmonates and other hormones (**Lager et al., 2010; Bailey et al., 2023**; **Jan et al., 2025; Singh et al., 2025**).

Therefore, pH_Ext_ is a central determinant of plant growth, development and stress responses, operating at the interface between the plant and its environment. It is known to influence root architecture through hormone-mediated pathways, regulate nutrient availability and uptake and integrates multiple signaling networks. However, its complexity and the lack of clearly defined sensing mechanisms have limited our understanding on how plants perceive and respond to the pH changes. Ongoing research efforts in identifying molecular mechanism of pH sensing using integration of multi-scale data are expected to advance this field with significant implications for both basic plant biology and agricultural applications. In our previous report we presented spatial abundance of auxin as the key factor in regulating root growth over a range of pH_Ext_ in Arabidopsis and that JA acts as the key upstream driver regulating the auxin accumulation in Arabidopsis roots (**Singh et al., 2025**), While no study has been carried out in crop plants so far, in the present work we have explored how rice responds against differential pH_Ext_. The study demonstrates that alike Arabidopsis, a highly conserved response against pH_Ext_ also operates in rice. In addition to this we have demonstrated that changing pH_Ext_; Fe availability in the media changes resulting into development of Fe toxicity and Fe deficiency responses towards acidic and alkaline pH conditions, respectively. Finally, the differential Fe toxicity/deficiency response act as the sensory module for pH_Ext_ and regulates jasmonate and auxin abundance in roots which in turn defines root growth inhibition.

## Material and Methods

### Plant growth and treatment conditions

Surface sterilization of rice (*Oryza sativa* var PB1) seeds was done with 0.1% mercuric chloride for 10 min followed by repetitive wash (5 times) with autoclaved MQ for 5 min each. Seeds were then kept on ½MS media with 3% sucrose and 0.8% agar for germination at 28°C for 4 days with 16/8h photoperiod. In order to assess the impact of pH_Ext_ on plant growth two strategies were opted i.e. hydroponics-based and soil-based. For hydroponics-based experimentation equally germinated seedlings were transferred to full strength liquid Yoshida media with custom-made floaters as done previously (**Singh et al., 2015**). Same number of equally germinated seedlings were then transferred in four liters of media for the indicated duration and conditions. Media was changed every 3^rd^ day while pH was maintained every day on same time. The concentration of nutrients in the media used was as follows: NH_4_NO_3_ (1.40mM), NaH_2_PO_4_ (0.32mM), K_2_SO_4_ (0.51mM), CaCl_2_.2H_2_O (1mM), MgSO_4_.7H_2_O (1.7mM), H_3_BO_3_ (19μM), ZnSO_4_.7H_2_O (0.15μM), CuSO_4_.5H_2_O (0.15μM), (NH4)_6_Mo_4_O_2_.4H_2_O (0.015μM), CitricAcid (70.75μM), Na-Fe-EDTA (60μM), and MnCl_2_.4H_2_O (9.46μM). Iron deficient media was achieved by excluding Na-Fe-EDTA from the media. MeJA stock (in DMSO, Sigma 392707-25ML) and NAA stock (in ethanol, Sigma N1641-50ML) were used for treatments with indicated dilutions. The experiments were conducted in a controlled growth chamber with 28°C, 16h/8h photoperiod with 280–300 μM photons/m2/s photon density and ∼70% humidity.

For soil-based experimentation, a mixture of soilrite:sand (3:1) was prepared and repetitively saturated with liquid Yoshida media with desired pH until the required pH was recorded. Equally germinated seedlings were planted to the indicated soil pH conditions (optimal growth conditions otherwise) and supplemented with liquid Yoshida media with respective pH for 30 days till the phenotype was recorded. The pH was checked every next day with fresh flow through media from the saturated soil when additional Yoshida media was supplemented.

Most of the phenotypic recording was done with a digital camera setup. For, WinRHIZO^TM^ imaging roots were counterstained with safranine. Lateral root length from WinRHIZO^TM^ images was quantified with ImageJ and data was plotted with Microsoft Excel with significance tests.

### Elemental detection

Elemental analysis in rice roots and shoots was performed using Inductively Coupled Plasma Mass Spectrometry (ICPMS) as mentioned previously (**Pandey et al., 2024**). Briefly, rice roots were thoroughly washed with a solution containing 2 mM CaSO₄ and 10 mM EDTA (pH 8.0) to remove any element adhered to the root surface. Following the wash, both root and shoot tissues were frozen in liquid nitrogen and ground to a fine powder using mortar and pestle. The powdered samples were then dried at 60 °C for 4–5 days until a constant dry weight was achieved. Once completely dried, the dry weight of each sample was recorded, and the material was transferred into acid-washed SuraPure™ PFA digestion tubes. Samples were digested with 5 ml concentrated nitric acid (HNO₃). The tubes were tightly sealed and placed in a microwave digestion system, where the temperature was gradually increased to 180 °C for complete digestion, following the instrument’s standard program. After digestion, the samples were diluted with 5% nitric acid to a known final volume. Calibration standards were prepared using certified multi-element standard solutions to generate a standard curve. Elemental concentrations in samples were recorded in three replicates and plotted accordingly.

### Ferrozine assay and Perl’s staining

To detect Fe^2+^ in the media ferrozine assay was conducted with the un-utilized media with different pH values where 0.3 mM ferrozine (working concentration) was added to the media followed by incubation for 1 hour in dark. The absorbance of the samples was taken at 562 nm using a plate reader and mean of the OD_562_ values for each pH conditions was plotted.

For colorimetric analysis of Fe assimilation in root tissues, Perl’s staining was done where freshly cut roots from the indicated treatments were washed and immerged in the preactivated solution containing 2% HCl and 2% potassium ferrocyanide. Pre-activation of the solution was done for at-least 30 minutes in dark at 37°C. The staining was allowed to be developed for 3 hours before washing with MQ water. Images were captured using stereo-zoom microscope.

### Transcriptome analysis

Rice root RNA was extracted after 10 days of different soil pH (pH_Ext_) treatments using RNeasy Plant Mini kit (Qiagen#74904) following the manufacturer’s instructions. Paired libraries were prepared from QC passed RNA samples using NEBNext^R^ Ultra^TM^ II Directional RNA library Prep Kit. PCR enriched libraries were purified using AMPureXP beads and quality of each library was checked before processing on Illumina platform. For each treatment three biological replicates were considered, and more than 10 million reads were obtained for each library. The resultant 09 libraries were mapped against the Oryza sativa genome (MSU7, http://rice.uga.edu/osa1r7_download/osa1_r7.asm.fa.gz) using STAR (v2.7.10a) with default parameters. FeatureCount (v2.0.3) and DEGSeq2 R package was used to perform differential gene expression analysis considering pH_5.8_ as the control conditions for both pH_3.8_ and pH_7.8_ treatments. For Principal component analysis (PCA) the RNA-seq data on phytohormone treatment in WT root tissues (along with replicates) were individually retrieved from Rice RNA-seq database (https://plantrnadb.com/ricerna/) (**Yu et al., 2022**). Libraries were manually selected and labelled for analysis. A total of 71 libraries were analyzed for auxin, abscisic acid (ABA), cytokinin (CK), ethylene (ET), jasmonic acid (JA), karrakin (kar) and strigalectone (Striga) treatment in WT root (**Table S1**). 19 libraries were analyzed for Fe deficient and 18 for high Fe treatment transcriptomic data retrieved from previously published datasets (**Table S2; Ueda et al., 2025**). Expression levels were normalized to their respective controls, and log_2_ fold change (Log_2_FC) values were used for PCA and heatmaps. PCA and heatmap images were generated using TB-tools (**Chen et al., 2020**). ShinyGO 0.85.1 (https://bioinformatics.sdstate.edu/go/) was used to perform GO enrichment analysis with all the significantly regulated genes (padj<0.05) on either pH_3.8_ or pH_7.8_, separately, considering *Oryza sativa* Japonica as the target databases with FDR cutoff (0.05).

#### qRT-PCR

Expression analysis with qRT-PCR was done as described previously (**Singh et al., 2021**). Briefly, total RNA was extracted from roots with TRIzol™ as per the manufacturer’s instructions followed by DNase I digestion. 1 μg of DNase I digested RNA was reverse-transcribed using High-Capacity cDNA Reverse Transcription Kit from Applied Biosystems™. qRT-PCR was performed on the Bio-Rad CFX96 Real-time PCR detection system using SYBR Green I (Takara SYBR Premix Ex Taq) with the gene specific primer sets (**Table S3**). The real-time PCR program was set as follows: 95°C for 3 min, 45 cycles (95°C for 10s, 55°C for 20s,72°C for 20s). All reactions were performed in three replicates for each data point. Fold change was calculated using 2^−ΔΔCT^ method, and Log_2_FC values were plotted. For normalization, the *Ubiquitin5* (Os01g0328400) was used as endogenous control (**Singh et al., 2020**).

#### Cell type cluster analysis

Genes expressing in different cell type clusters were taken from previously published root specific single-cell-RNAseq data (**Liu et al., 2021**). Expression of all the genes in different cell type clusters was analyzed and only significantly and differentially expressed genes were considered for plotting. Violin plots with differential expression of genes associated cell types was plotted with R using ggplot2 package.

#### Cross-species validation

For the cross-species analysis of transcriptomic behavior in rice and Arabidopsis, the list associated with gene-gene orthologues in rice and Arabidopsis was taken from the previous study (**Liu et al., 2021**). Expression patten of differentially regulated genes in Arabidopsis for acidic and alkaline pH was taken from previously published expression datasets (**Lager et al., 2010, Bailey et al., 2023**). All the genes were mapped with rice transcriptome datasets manually and dot-plot was generated using Microsoft Excel with genes commonly and significantly (Log2FC≥0.5; padj<0.05) expressed in both rice and Arabidopsis.

### Hormonal quantification

Phytohormone quantification was done as per our previous study (**Singh et al., 2020; Singh et al., 2021**) with minor modifications. Briefly, whole root tissues were harvested and immediately frozen in liquid nitrogen and stored at -80 °C until used. Approximately, 200mg tissue was pulverized and processed in 1 ml cold methanol with the addition of internal standards (D_6_-JA-Ile or D_5_-IAA). The homogenate was vortexed for 30 min followed by centrifugation at 14,000 rpm for 20 min at 4°C. Quantification was done in five replicates using Exion LC coupled to Triple Quad 6500+ (Sciex).

## Results

### Arabidopsis and rice show conserved response against varying rhizospheric pH conditions

Our previous report (**Singh et al., 2025**), demonstrated that any deviation in pH_Ext_ produces an inhibitory effect on Arabidopsis root growth. To explore how crop plants like rice with fibrous root system architecture (RSA) responds to the varying pH_Ext_ conditions, rice seedlings were subjected to different pH_Ext_ conditions [pH_3.8_ (acidic), pH_5.8_ (optimal) and pH_7.8_ (alkaline)] and compared the RSA parameters. Our observations marked similar responses of root growth towards different rhizospheric pH conditions in both the plant species. Notably, rice root growth was significantly inhibited towards both acidic and alkaline pH_Ext_ conditions alike Arabidopsis (**Figure 1A-D**). Interestingly, shoot growth doesn’t show significant inhibition on alkaline pH_Ext_ though inhibition was observed in terms of length and dry weight on acidic pH_Ext_ conditions (**Figure 1E-G**). The root length here (**Figure 1B**) corresponded to the longest root where it excluded the variation among seminal root length within the plant, to overcome this limitation, we analyzed the finer details of RSA using WinRHIZO^TM^ (https://regent.qc.ca/assets/winrhizo_about.html) image acquisition system. Our results show a typical root growth inhibition (RGI) response towards both acidic as well as alkaline rhizospheric pH conditions affecting the average seminal root length (**Figure 1H-I**). The finer observations on RSA further revealed increase in lateral root length on increasing rhizospheric pH **(Figure 1J)**.

**Figure 1.**
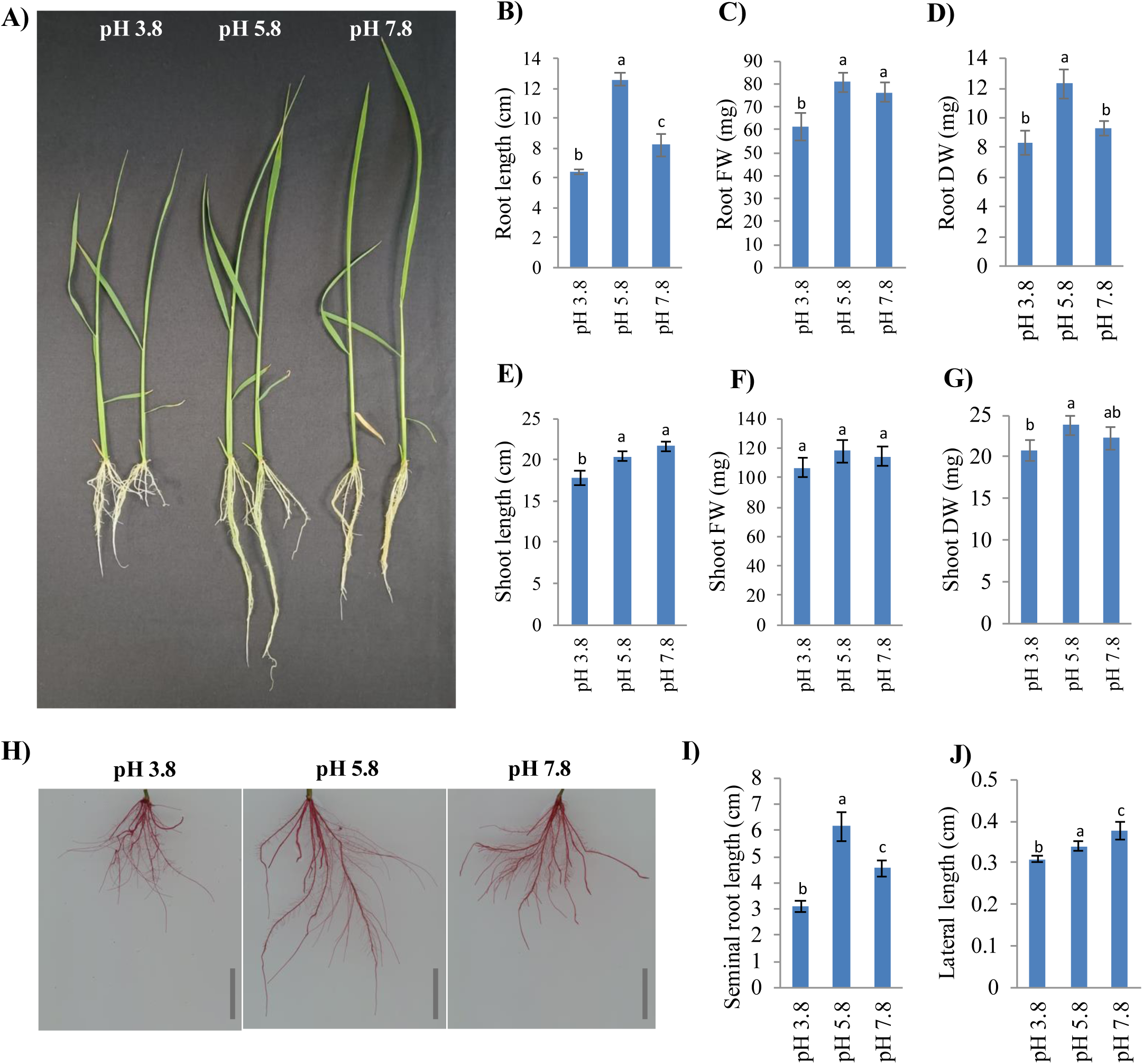
Rhizospheric pH severely impacts growth and root system architecture in rice. Uniformly germinated rice seedlings were transferred to Yoshida liquid media with mentioned pH conditions. Growth parameters were analyzed after 10 days of treatment under mentioned condition. **A)** Representative seedling growth under respective growth condition, **B-D)** Quantitative analysis of root growth parameters (n≥10), **E-G)** Shoot growth parameters (n≥10), **H)** Representative images showing root system architecture under different growth conditions, Quantitative analysis of average **I)** seminal root length (5 plants) and **J)** lateral root length (n³50) under different rhizospheric pH conditions. Mean values were plotted with error bar representing SE among the replicates and significance was analyzed using ANOVA, groups with different letters differ significantly based on Benjamini–Hochberg-adjusted post hoc pairwise *t*-tests (*P* < 0.05).

Next, we analyzed the pattern of root growth under soil system with different pH_Ext_ conditions. After 30 days of treatment, we observed a severe inhibition in root and shoot growth on both the pH_Ext_ extremes. Notably, ∼50% growth inhibition was observed in root length and ∼95% inhibition was observed in root FW and DW on pH_3.8_ (**Figure 2A-E**). The root also showed inhibition on alkaline pH_Ext_ conditions although the level of inhibition was less pronounced but significant i.e. root length, FW and DW, ∼20%, ∼36% and ∼26% inhibition, respectively. Besides root, shoot growth also showed inhibition on both the pH extremes i.e. ∼35% and ∼34% in shoot length (**Figure 2F**), ∼86% and ∼30% in shoot FW and ∼91% and ∼39% in shoot DW on pH_3.8_ and pH_7.8_, respectively when compared to pH_5.8_ (**Figure 2G-H**). These observations suggested that rhizospheric pH has ability to severely impact root system architecture and that any deviation from optimal pH would lead to root growth inhibition in rice while increasing pH_Ext_ may have stimulatory effects on lateral root elongation.

**Figure 2.**
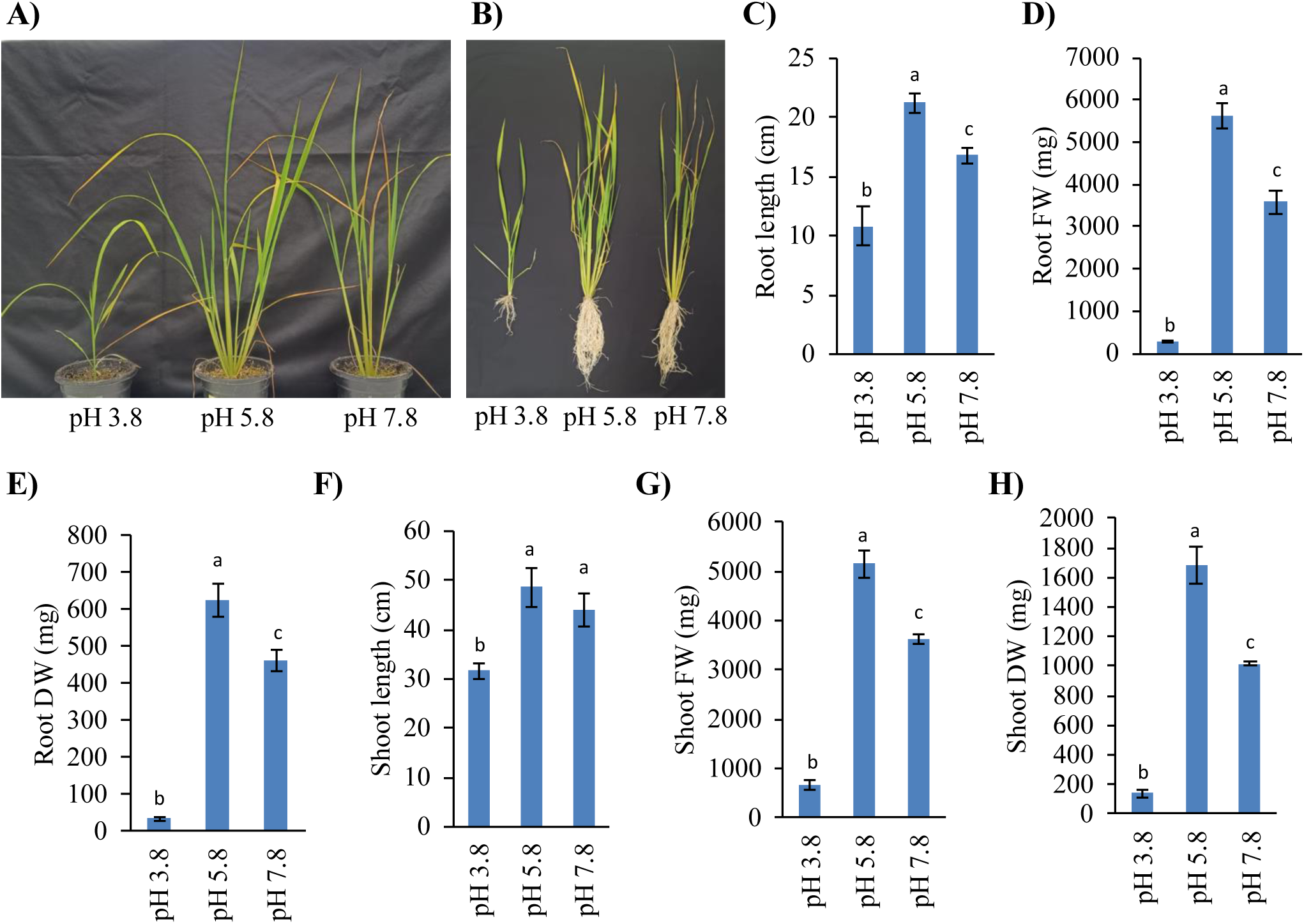
Effect of rhizospheric pH on rice growth in soil conditions. Uniformly germinated rice seedlings were transferred to soilrite:sand (3:1) mixture (saturated with Yoshida liquid media with mentioned pH until no further change was observed in pH). Plants were analyzed after 30 days of growth. **A)** Plant growth in pots, **B)** uprooted plants showring root growth, **C)** average root length, **D)** root fresh weight, **E)** root dry weight, **F)** shoot length, **G)** shoot fresh weight and **H)** shoot dry weight. Each datapoint represent mean of three biological replicates with error bar representing SE among the replicates. Significance was analyzed using ANOVA, groups with different letters differ significantly based on Benjamini–Hochberg-adjusted post hoc pairwise *t*-tests (*P* < 0.05).

### Transcriptomic response of rice roots against rhizospheric pH

To elucidate the mechanism behind the phenotypic observations against rhizospheric pH, we conducted root specific mRNA sequencing (RNA-seq) from rice seedlings subjected to different pH_Ext_ conditions. A total of 6358 genes were found differentially regulated (Log_2_FC>0.5, Padj<0.05), on either of the treatment conditions out of which 1091 genes were commonly influenced in both the treatments (**Figure 3A**). Further, 3273 genes were differentially regulated on pH_3.8_ out of which 2206 genes were upregulated, and 1067 genes were downregulated. Whereas 4176 genes were differentially regulated on pH_7.8_ treatment out of which 2040 genes were upregulated, and 2136 genes were downregulated. The comparative analysis of differentially regulated genes suggested that 2182 genes were exclusively responsive to pH_3.8_ when compared to pH_5.8_; whilst 3085 genes were exclusively responsive to pH_7.8_. To our interest, 536 genes were found to have opposite expression behavior to different pH_Ext_ extremes, among them, 399 genes were found upregulated on pH_3.8_ and downregulated on pH_7.8_ in contrast, 137 were found upregulated on pH_7.8_ and downregulated on pH_3.8_. In addition to this, 374 genes were found upregulated and 181 genes were found downregulated to both the condition. While the genes showing inverse expression pattern across the pH_Ext_ range may be linked closely to the response against changing pH, the genes commonly upregulated or downregulated may be correlated to the root growth behavior.

**Figure 3.**
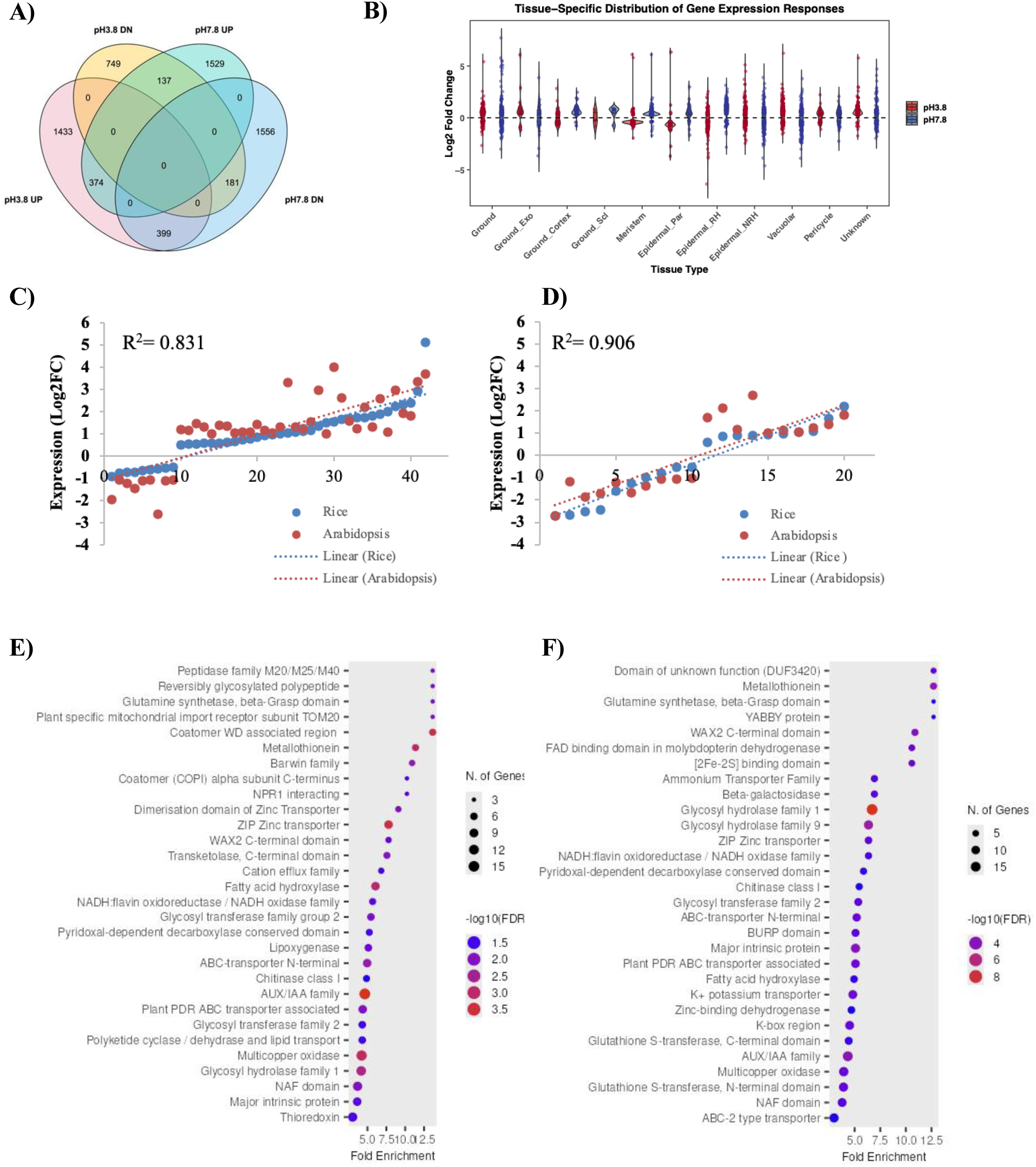
Transcriptomic landscape of rice and Arabidopsis roots in response to acidic and alkaline pH conditions. **A)** Venn diagram indicating number of differentially regulated genes in rice roots under different pH conditions **B)** Violin plot showing transcriptional behavior of differentially regulated genes preferentially expressing in different cell types of rice root. **C-D)** Dot plots showing correlation among the differentially expressed genes in rice and Arabidopsis in response to acidic and alkaline pH conditions, respectively. **E-F**) GO enrichment analysis of genes under acidic and alkaline pH conditions, respectively.

Now, having root specific transcriptome, we wanted to explore if rhizospheric pH has any specific influence on certain distinct cell types in roots. Therefore, we analyzed how the rhizospheric pH changes the expression pattern of genes preferentially expressed in different cell type clusters. For this, we utilized previously defined cell type clusters identified in rice root tissues using single-cell RNA-seq (**Liu et al., 2021**) analysis and observed how the expression pattern of these clusters is regulated on changing pH_Ext_ (**Figure 3B**). The analysis suggested changes in almost every cell type clusters but considerable changes were observed in ground_exodermis (ground_exo), ground_cortex as well as ground_sclerenchyma (ground_scl). Further, the observations also marked notable changes in root meristem and epidermis specifically epidermal_parental (epidermal_par) and epidermal_root hair (epidermal_RH) cells. Therefore, the analysis suggested a broad scale changes in the transcriptomic pattern of genes in roots subjected to differential pH conditions.

Next, as the phenotypic response against rhizospheric pH was observed to be highly conserved among rice and Arabidopsis, we wanted to explore if the response is also conserved at the transcriptomic level. For this, talking advantage of already identified orthologous genes between the two species (**Liu et al., 2021**), we performed correlation analysis between our dataset in rice and the differentially expressed genes reported in Arabidopsis (**Lager et al., 2010; Bailey et al., 2023**). Considering common genes identified in both the species with significant change in expression, we observed that despite of the anatomical and developmental differences, both the species showed a very similar transcriptomic response against rhizospheric pH with very high Pearson correlation coefficient (R^2^=0.831 and R^2^=0.906 during acidic and alkaline pH, respectively) (**Figure 3C-D**). After having established the conserved phenotypic and transcriptomic response among rice and Arabidopsis, we conducted GO enrichment analysis highlighting conserved protein families in the differentially regulated gene sets. The GO enrichment highlighted photosynthesis, glutamine synthesis, zinc transporters, fatty acid hydroxylation, cation export, lipoxygenase, auxin response, glutathione-S-transferase, iron homeostasis and redox reactions as the key pathways commonly targeted by the acidic and alkaline pH in rice roots (**Figure 3E-F**). Therefore, the transcriptome analysis in rice and Arabidopsis while projected a conserved response against rhizospheric pH among plant species, the enrichment of similar pathways under acidic and alkaline pH treatments further suggested rhizospheric pH as a continuous external stimulus for the plants.

### Auxin and JA crosstalk explains pH_Ext_ mediated RGI in rice

As established in our previous report, auxin and JA signaling crosstalk at root tip plays a pivotal role in pH_Ext_ mediated RGI in Arabidopsis (**Singh et al., 2025**). The transcriptome analysis in Arabidopsis also highlighted enrichment of multiple hormone responsive genes (**Lager et al., 2010; Bailey et al., 2023**). A recent report in rice further suggested involvement of ABA in the low pH tolerance (**Jan et al., 2025**). These reports along with the GO terms enrichment of auxin and JA signaling (lipoxygenase/fatty acid hydroxylase) in our transcriptome data projected a possible involvement of multiple hormones in plant’s response to pH_Ext_. To investigate this aspect, we compared transcriptomic responses of rice root under different phytohormone treatment conditions with the one associated with acidic (pH_3.8_) and alkaline pH (pH_7.8_) conditions from our transcriptome dataset. The PCA analysis suggested that the transcriptomic changes due to rhizospheric pH (both acidic and alkaline) were overlapping to that observed on auxin, cytokinin (CK), ethylene (ET), strigolactone (striga) and karrakin (kar) treatments (**Figure 4A**). Interestingly, JA and ABA treatment forms two separate clusters indicating no overlap with the pH mediated transcriptomic responses. Considering that a larger portion of the transcriptome may not be related to any of the phytohormone and there is a possibility of crosstalk among the phytohormones which might not be reflected in this analysis, we investigated individual hormone signaling genes under the given pH_Ext_ conditions. The analysis revealed representation of all major phytohormone signaling genes in the transcriptome data but the enrichment of jasmonate and auxin was specifically notable (**Figure 4B**). Induction of JA, ABA and SA signaling/biosynthesis genes was notable on pH_3.8_ with downregulation of these genes on pH_7.8._ Exceptionally, auxin showed majority of genes induced on pH_7.8_ and either downregulation or no change on pH_3.8_. These observations were in justification to our previous report where JA and auxin signaling were demonstrated to have inverse pattern and antagonistic crosstalk on varying pH_Ext_ treatments in Arabidopsis (**Singh et al., 2025**). While ABA signaling was reported to be involved in plants response to low pH, its induction at pH_3.8_ was justifiable (**Jan et al., 2025**). Having known that higher intrinsic JA content/signaling may induce ABA biosynthesis in rice (**Singh et al., 2021**), and similarly, involvement of SA in multiple biotic and abiotic responses (**Tian et al., 2025**) led us to speculate that ABA and SA may have regulatory roles in plant’s response to pH_Ext_.

**Figure 4:**
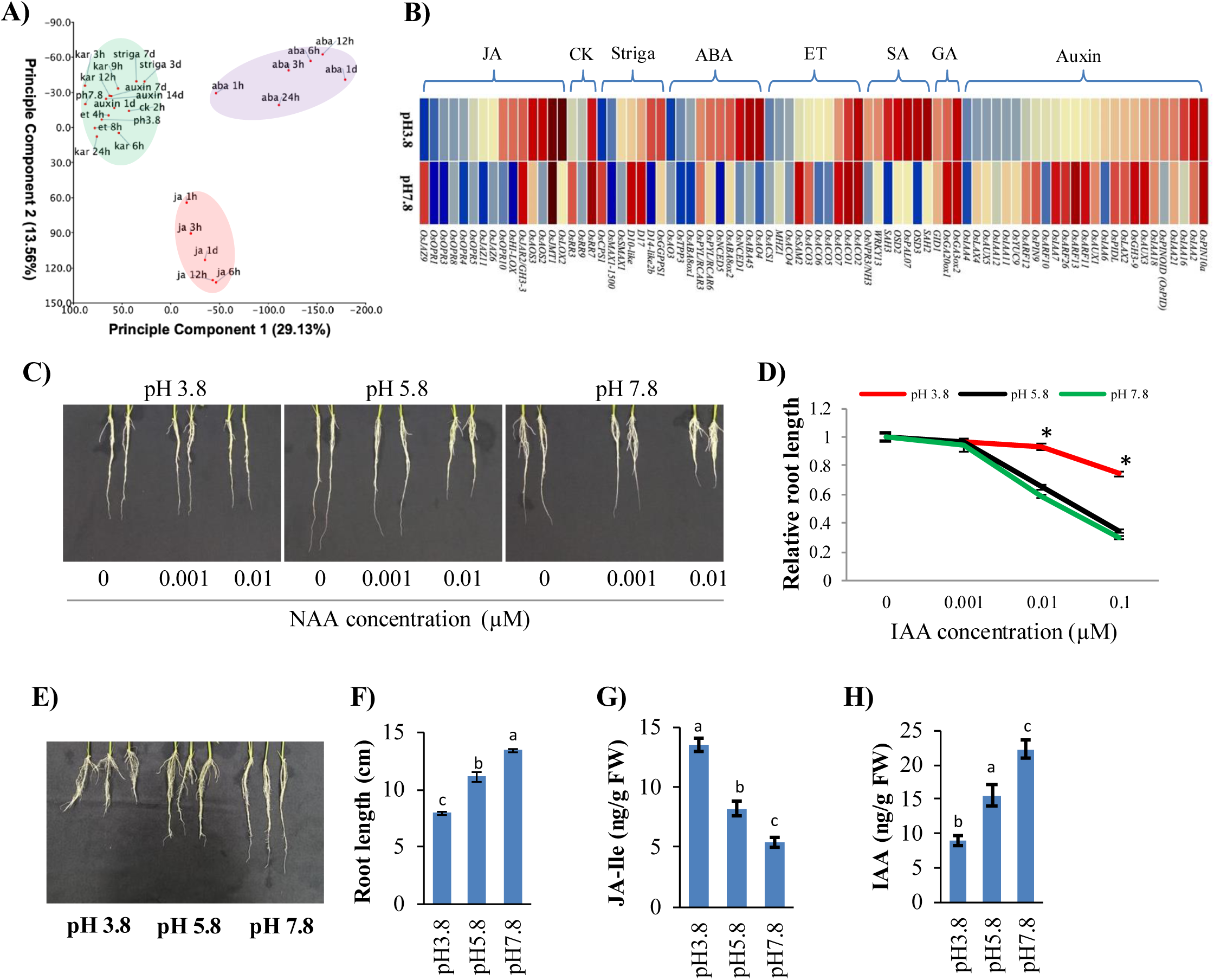
Hormonal regulation of rice response against rhizospheric pH. **A)** PCA analysis showing overlapping transcriptional response of rice roots against rhizospheric pH and different phytohormone treatments (auxin, JA, ABA, cytokinin (CK), ethylene (ET), strigolactone (striga) and karrakin (kar)). **B)** Heatmaps showing expression pattern of hormone specific/signaling genes in rice roots against pH_3.8_ and pH_7.8_ as compared to pH_5.8_. Log_2_FC values were plotted. **C-D)** Effect of NAA treatment (indicated concentrations) on rhizospheric pH mediated RGI after 10 days of treatment (n≥15). Relative root length was calculated for different NAA concentrations by dividing it by root length at 0µM NAA treatment for the particular pH treatment. **E-F)** Effect of MeJA (0.01µM) treatment on rhizospheric pH mediated RGI after 10 days of treatment (n≥15). **G)** JA-Ile (n=5) and **H)** auxin content (n=5) in rice root after 10 days of growth on different rhizospheric pH conditions. Bars indicate the mean of the biological replicates with error bars representing SE among the replicates. Significance was analyzed using ANOVA, groups with different letters differ significantly based on Benjamini–Hochberg-adjusted post hoc pairwise *t*-tests (*P* < 0.05).

In order to delineate the role of auxin and JA in pH_Ext_ mediated RGI, we analyzed root growth of rice seedlings subjected to different pH_Ext_ conditions with either 1-Naphthaleneacetic acid (NAA) or methyl-jasmonate (MeJA) treatment. Expectedly, rice seedlings with NAA treatment were hyposensitive to pH_3.8_ mediated RGI compared to pH_5.8_ while no significant difference in root growth rate was observed on pH_7.8_ though it was relatively hypersensitive (**Figure 4C-D**). In contrast, when seedlings were subjected to different pH conditions along with MeJA, root growth behaved insensitive to pH_7.8_ and hypersensitive to pH_3.8_ mediated RGI (**Figure 4E-F**). These experiments confirmed the involvement of auxin and JA signaling in pH_Ext_ mediated RGI and that too in different pH_Ext_ extremes, as observed in Arabidopsis. To further validate these observations, we quantified JA and auxin levels in rice roots subjected to different pH_Ext_ conditions for 7 days. Our observations revealed decreasing accumulation of JA-Ile (**Figure 4G**) and simultaneously, increasing accumulation of IAA (**Figure 4H**) in root tissues on increasing pH_Ext_. These observations not only justified the transcriptomic pattern shown by JA and auxin responsive/signaling genes (**Figure 4B**) but also the rescue of pH_3.8_ mediated RGI on NAA treatment and the rescue of pH_7.8_ mediated RGI on MeJA treatment. Therefore, the RGI on pH_3.8_ strongly corresponds to high JA and low auxin accumulation whereas the RGI on pH_7.8_ corresponds to low JA and high auxin accumulation in roots.

Next, to comment whether JA and auxin signaling interacts with each other, we analyzed how auxin treatment influence JA signaling/biosynthesis genes and conversely, how JA treatment influence auxin related genes. For this, we extracted previously published root specific transcriptome datasets on auxin and JA treatment and re-analyzed all the significantly influenced auxin or JA related genes. The observations revealed only a weak negative influence of auxin treatment on JA signaling genes with slight induction of JA signaling repressors (*OsJAZs*) and downregulation of JA biosynthetic genes (*OsAOS4*, *OsOPR9*, *OsOPR10*, *OsOPR6*). Further, the JA associated transcriptome was found significantly induced on JA treatment validating the concept that JA signaling genes are inducible to JA treatment (**Figure S1A; Wasternack and Hause, 2013**). Interestingly, when auxin signaling genes were analyzed on JA treatment, a prominent downregulation of auxin signaling and biosynthetic genes was observed (**Figure S1B**). Therefore, while auxin may not have a strong influence on JA signaling, JA seems to act antagonistically to auxin signaling. Considering the remarkable antagonistic regulation, increasing auxin accumulation on increasing pH_Ext_ can therefore be speculated as the consequence of decreasing JA accumulation. Also, as reported in Arabidopsis, the observations strengthened the notion that rhizospheric pH regulate JA accumulation which in turn regulate auxin to define RGI pattern, as a conserved response in rice and Arabidopsis.

### Rhizospheric pH mediated RGI correlates with Fe accumulation in root

Rhizospheric pH is well known to influence nutrient availability, and it is often considered that plant experiences several micronutrient fluctuations and heavy metal toxicities which may regulate plant’s response to a particular rhizospheric pH (**Tsai and Schmidt, 2020; Tsai and Schmidt, 2021**). To understand if rhizospheric pH has any influence on nutrient status of plants, we analyzed the micronutrient assimilation in root and shoot tissues of rice seedlings subjected to different pH_Ext_ conditions. The ICPMS analysis revealed increasing assimilation of micronutrients like Zn, Mn, Mg, Co and Cu in rice roots on increasing rhizospheric pH values (**Figure 5A**). Also, this increase in root micronutrient assimilation is well replicated in shoot tissues except for Mg and Cu, which showed no significant change in shoot tissues due to pH_Ext_ change. Interestingly, Fe accumulation in root as well as shoot tissues showed an inverse pattern compared to other micronutrients where decreasing Fe assimilation was observed on increasing rhizospheric pH. Therefore, to study the Fe responses on differential pH conditions, we analyzed how Fe responsive genes were influenced on varying pH_Ext_. The analysis of our transcriptome data revealed exceptionally high induction of Fe uptake machinery on pH_7.8_ indicating a severe Fe deficient condition (**Figure 5B**). While the induction of Fe deficiency response on pH_7.8_ was justifiable considering the reduced Fe assimilation in root (0.60-fold) and shoot (0.63-fold); the observed non-significant change in Fe deficiency responsive transcriptome on pH_3.8_ even after accumulating higher Fe in root (1.86-fold) and shoot (1.67-fold) was surprising. Although, induction of key genes including *OsVIT2*, *OsFER1-2* and downregulation of *OsMIR*, *OsIRT1* and *OsNRAMP1* favors Fe toxicity response on pH_3.8_ (**Figure 5B; Mahender et al., 2019; Aung and Masuda, 2020**). This ambiguity of Fe responses against differential pH_Ext_ made us to investigate the RGI pattern considering the Fe availability and pH_Ext_ simultaneously. For this we subjected rice seedlings to media with different pH_Ext_ along with Fe sufficient or Fe deficient conditions. While considering only optimal pH (pH_5.8_) during Fe sufficient and Fe deficient conditions, root length inhibition on Fe deficiency validated the typical Fe deficiency response, and therefore our experimental conditions (**Figure 5C-D**). Interestingly, Fe deficient condition completely rescued pH_3.8_ mediated RGI and produced hypersensitive RGI response on pH_7.8_ (**Figure 5C-D**). To further investigate this behavior, we analyzed Fe accumulation using Perl’s stain and ICPMS under Fe sufficient and Fe deficient conditions (**Figure 5E-G**). Both, intensity of the staining and ICPMS data points towards the hyperaccumulation of Fe on pH_3.8_ and acute Fe deficiency on pH_7.8_ with a general trend where Fe accumulation in root and shoot tissues decreases on increasing rhizospheric pH values. The phenotypic observations along with Fe content in root and shoot, all suggested that Fe can be the critical determinant of root growth inhibition on varying pH_Ext_ conditions, where, both Fe toxicity and deficiency are known to inhibit root growth (**Fageria et al., 2008; Sun et al., 2017**).

**Figure 5.**
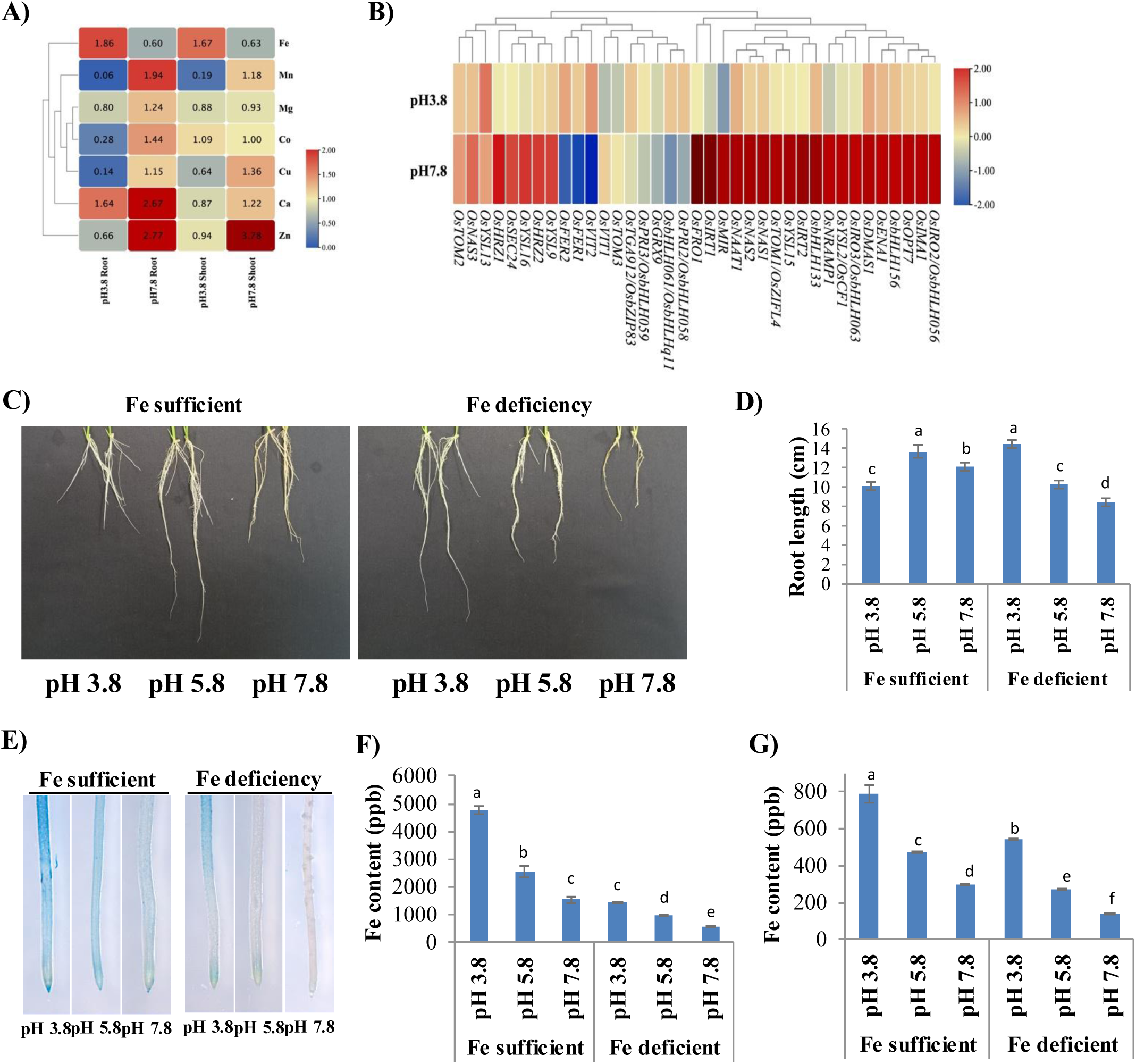
Rhizospheric pH mediated RGI correlates with Fe accumulation in root. **A)** Change in micronutrient assimilation (fold change) in rice root and shoot after 10 days of growth on different rhizospheric pH as observed with ICPMS analysis. Fold change was calculated with reference to pH_5.8_. **B)** Heatmap showing expression pattern of Fe deficiency responsive genes on pH_3.8_ and pH_7.8_ as compared to pH_5.8_ in the root specific transcriptome analysis. Log_2_FC values were plotted. All the genes which showed significant change in either of the treatment were considered. **C-D)** Representative images and quantitative analysis showing effect of rhizospheric pH on root length under Fe sufficient and deficient conditions after 10 days of treatment (n³10). **E)** Perl’s staining showing Fe assimilation in rice root under different conditions. **F-G)** ICPMS analysis showing Fe assimilation in root and shoot tissues, respectively, after the treatment. Mean values were ploptted with error bar representing SE among the replicates and significance was analyzed using ANOVA, groups with different letters differ significantly based on Benjamini– Hochberg-adjusted post hoc pairwise *t*-tests (*P* < 0.05).

### Fe deficiency response participates in rhizospheric pH perception in rice

Our results point that JA, auxin and Fe responsive genes participate in pH_Ext_ mediated RGI. Next, we investigated if Fe response work independently or interacts with JA-auxin crosstalk to regulate pH_Ext_ mediated RGI? For this, we analyzed effect of Fe toxicity and deficiency on auxin and JA associated genes. For this we again utilized previously published multiple root specific RNA-seq datasets under Fe toxicity and Fe deficient conditions from the perspective of auxin and JA signaling/biosynthesis genes (**Figure 6A**). The analysis revealed notable induction of JA signaling/biosynthesis genes and inhibition of auxin related transcriptome on Fe toxicity. Fe toxicity prominently repressed auxin biosynthesis and signaling with induction of known auxin catabolism genes like *OsGH3-2* **(Du et al., 2012)**. When similar analysis was conducted on Fe deficiency treatment the high expression of multiple auxin biosynthesis (*OsTAR2* & *OsYUC9*) and signaling (Os*AUX1/OsLAX1*, Os*AUX3/OsLAX3*, Os*ARF13, OsIAA25)* genes was observed. In addition to this, known JA biosynthesis genes including *OsOPR1*, *OsOPR3*, *OsOPR6*, *OsOPR10*, *OsHI-LOX* were found downregulated on Fe deficiency treatment. As this analysis was conducted with previously published datasets, we validated the expression pattern of JA and auxin related genes under Fe sufficient and Fe deficient condition over the varying pH in our experimental setup (**Figure S2**). The qRT-PCR based analysis validated our transcriptomic observations under control condition (Fe sufficient) with very high degree of correlation. Also, an induced auxin signaling and inhibited JA signaling was observed on Fe deficient condition suggesting that the two signaling pathways are inversely regulated by Fe deficiency response. Now considering that the acidic pH_Ext_ mimicking Fe toxicity and alkaline pH mimicking Fe deficient conditions, the observed high JA-Ile and low auxin accumulation on acidic pH as well as low JA-Ile and high auxin accumulation on alkaline pH (**Figure 4G-H**) both are well justified as the function downstream of Fe toxicity/deficiency responses (**Figure 6A**).

**Figure 6.**
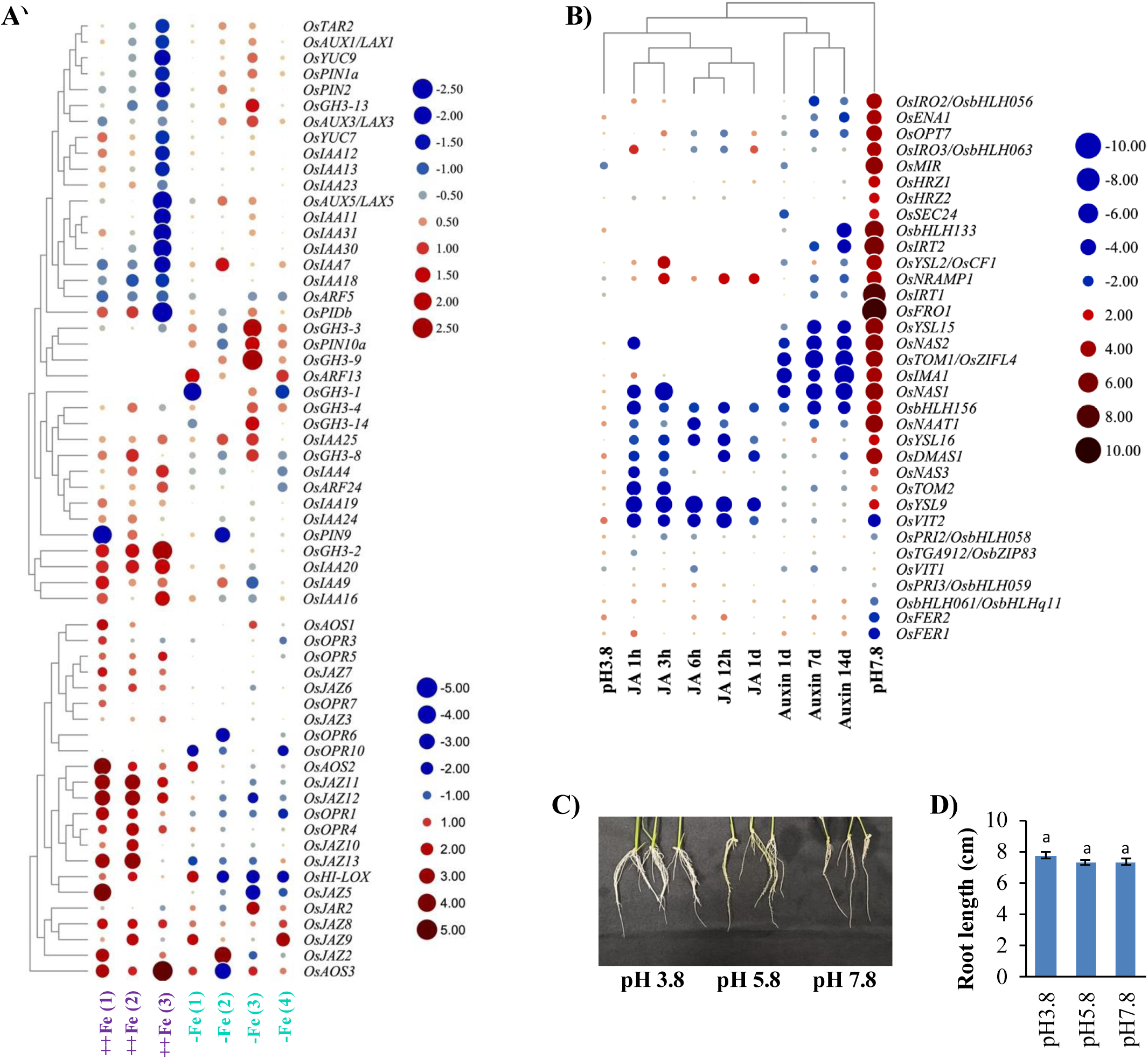
Iron toxicity and deficiency differentially regulates auxin or JA signaling. **A)** Expression pattern of auxin and JA signaling/biosynthesis genes in rice roots under Fe toxicity and deficiency conditions. **B)** Heatmap showing effect of JA and auxin treatment on Fe deficiency responsive genes. Log_2_FC values were plotted with genes having expression Log_2_FC >0.5 in any of the RNA-seq data. **C-D)** Representative images and root length quantification showing effect of varying rhizospheric pH along with MeJA (0.01µM) and Fe deficiency treatment, simultaneously, after 10 days of treatment (n≥15). Mean values were plotted with error bar representing SE among the replicates. Significance was analyzed using ANOVA, groups with different letters differ significantly based on Benjamini–Hochberg-adjusted post hoc pairwise *t*-tests (*P* < 0.05).

To further validate this opinion, we analyzed transcriptional behavior of Fe deficiency related genes on auxin and JA treatment which suggested downregulation of major Fe deficiency response gene on either JA or auxin treatments. Interestingly, entirely different set of Fe deficiency response genes were downregulated by either of the treatment. Genes such as *OsIMA1, OsTOM1/OsZIFL4, OsNAS1, OsNAS2, OsbHLH156, YSL15, IRT2, OsbHLH133, IRT1, ENA1* and *OsIRO2/OsbHLH056* were down regulated, on auxin treatment, whereas *OsNAS2, OsNAS1, OsYSL9, OsTOM2, OsNAS3, OsDMAS1, OsVIT2, OsYSL16, OsbHLH156* and *OsNAAT1* were downregulated on JA treatment (**Figure 6B**). These observations resulted into the conflict where the transcriptomic pattern on alkaline pH conditions despite of having higher auxin accumulation showed induced Fe deficiency response. Similarly, JA treatment also inhibits Fe deficiency transcriptome, but pH_3.8_ treatment exhibiting higher JA do not have a clear downregulated Fe deficiency response. In other words, neither higher JA during acidic pH nor auxin during alkaline pH complements the Fe deficiency response observed during pH_3.8_ and pH_7.8_ conditions, respectively. Therefore, the Fe deficiency response seems not to be regulated downstream of auxin and/or JA signaling, instead, JA and auxin signaling can be well correlated downstream of Fe assimilation; indicating that Fe deficiency response work upstream of JA-auxin crosstalk. The observations that JA treatment rescued pH_7.8_ mediated RGI (**Figure 4E-F**) without complementing the Fe accumulation in root **(Figure 5F; Figure S3A)** further validated the hypothesis that pH_7.8_ mediated RGI is regulated via JA deficiency or auxin toxicity in root and that Fe may work upstream of JA/auxin signaling during the process. To test this hypothesis, we extended our observations with MeJA treatment and analyzed if Fe deficiency can restore the rescued alkaline pH (pH_7.8_) mediated RGI as observed on MeJA treatment (**Figure 4E-F**). Here our experiment revealed that creating Fe deficient conditions completely abolished MeJA mediated rescue of RGI on pH_7.8_ (**Figure 6C-D**). While Perl’s stain confirmed Fe deficiency in rice roots (**Figure S3B**), the rescue of pH_3.8_ mediated RGI by Fe deficiency (**Figure 5C-D**) was also abolished (**Figure 6C-D**) on MeJA treatment. These observations strongly indicated that Fe deficiency mediated rescue of RGI on pH_3.8_ (**Figure 5C-D**) was facilitated via suppression of JA signaling (**Figure 6A**). Therefore, when exogenous MeJA was applied, Fe deficiency failed to nullify the effect of MeJA treatment sufficiently resulting into the restored RGI (**Figure 6C-D**). On the other hand, failure of MeJA treatment to rescue pH_7.8_ mediated RGI during Fe deficient conditions (**Figure 6C-D**) can also be justified considering that while pH_7.8_ itself mimics Fe deficient condition, creating further Fe deficiency strongly inhibit JA signaling (**Figure 6A**) leaving it inefficient for RGI rescue. The results can strongly be speculated that Fe toxicity at acidic pH conditions results into induced JA signaling, on the other hand, when pH increases, Fe deficiency is established inhibiting JA signaling towards alkaline pH. As JA antagonistically regulate auxin signaling, auxin signaling follows inverse pattern to what observed with JA. Therefore, Fe deficiency response seems to work as the sensory module which may regulate both JA and auxin towards both the extremes of pH_Ext_ and the final dialogues between JA and auxin defines the RGI pattern over the entire range of pH_Ext_.

### Fe bio-availability decreases on increasing rhizospheric pH

Considering the ICPMS data (**Figure 5F-G**), we knew that increasing rhizospheric pH develops Fe deficiency in rice. Next, we wanted to explore how Fe deficiency is developed on increasing pH_Ext_ despite of media being supplemented with equal Fe-EDTA. For this we conducted ferrozine assay with un-utilized media which revealed consistent decrease in the Fe^2+^ availability in the media on increasing pH (R^2^= -0.9526) (**Figure 7A-B**). These observations agreed with the previous reports where decreased availability of Fe^2+^, Fe^3+^, Fe(OH)_4_^+^ and increased availability of Fe(OH)_4_^-^ was observed on increasing rhizospheric pH (**Marschner, 2012**). Knowing that rice also use strategy-II for Fe acquisition where phytosiderophores (PS) are released into the rhizosphere which binds Fe^3+^ and the complex (Fe^3+^-PS) is absorbed by the roots (**Wairich et al., 2019**). Induction of all the genes associated with PS synthesis (*OsNAS1, OsNAS2, OsNAS3, OsNAAT1, OsDMAS1*), efflux (*OsTOM1, OsTOM2, OsTOM3, OsENA1*) and uptake (*OsYSLs*) though suggested activation of strategy-II for Fe uptake on pH_7.8_ but failed to justify the Fe deficient status of the plants (**Figure 5B**, **Figure 5F-G**). To comment on whether strategy-II in rice is functional at alkaline pH, we revisited ICPMS data and observed that while Fe content show decreased assimilation in root (0.60-fold) and shoot (0.63-fold) on pH_7.8_, Zn accumulation was strikingly higher in root (2.77-fold) as well as in shoot (3.78-fold) (**Figure 5A**). Knowing that Fe acquisition system can also facilitate Zn uptake, we speculated that the induced strategy-II facilitated Zn^2+^ uptake because the media was highly deficient in Fe content. It is known that at alkaline pH, Zn remain comparatively more soluble consequently, the higher availability of Zn competitively favors Zn^2+^-PS complex formation over Fe^3+^-PS complex (**Marshner, 2012; Schenkeveld et al., 2014**; **von Wiren et al., 1996**). The hypothesis can further be validated by analyzing correlation between Fe and Zn content in root (R^2^= -0.8357) and shoot (R^2^= -0.7828) tissues on increasing rhizospheric pH **(Figure S4A-B**). The correlation further improved for root (R^2^= -0.9787) and shoot (R^2^= -0.9664) during Fe deficiency because competition from Fe was further reduced (**Figure S4C-D**). Therefore, these observations strongly suggest that strategy-II which was meant for Fe acquisition become inefficient during alkaline pH where competitive acquisition of Zn dominates Fe acquisition leaving plants Fe deficient. The Fe deficient behavior of rice seedlings despite of induction of strategy-II can further be justified knowing that the Fe availability on alkaline pH (pH_7.0_ to pH_9.0_) is too low (∼10^-10^M) than required for the optimal growth (10^-6^ to 10^-5^ M) (**Boukhalfa and Crumbliss, 2002; Kraemer, 2004; Marschner, 2012**). Therefore, the results can be summarized as the increasing pH_Ext_ consistently reduces Fe availability in the rhizosphere which is differentially deduced by the plants in the form of relative Fe toxicity or Fe deficiency which in turn regulate JA-auxin crosstalk resulting into the observed RGI on either side of the optimal pH_Ext_ (**Figure 7C**).

**Figure 7.**
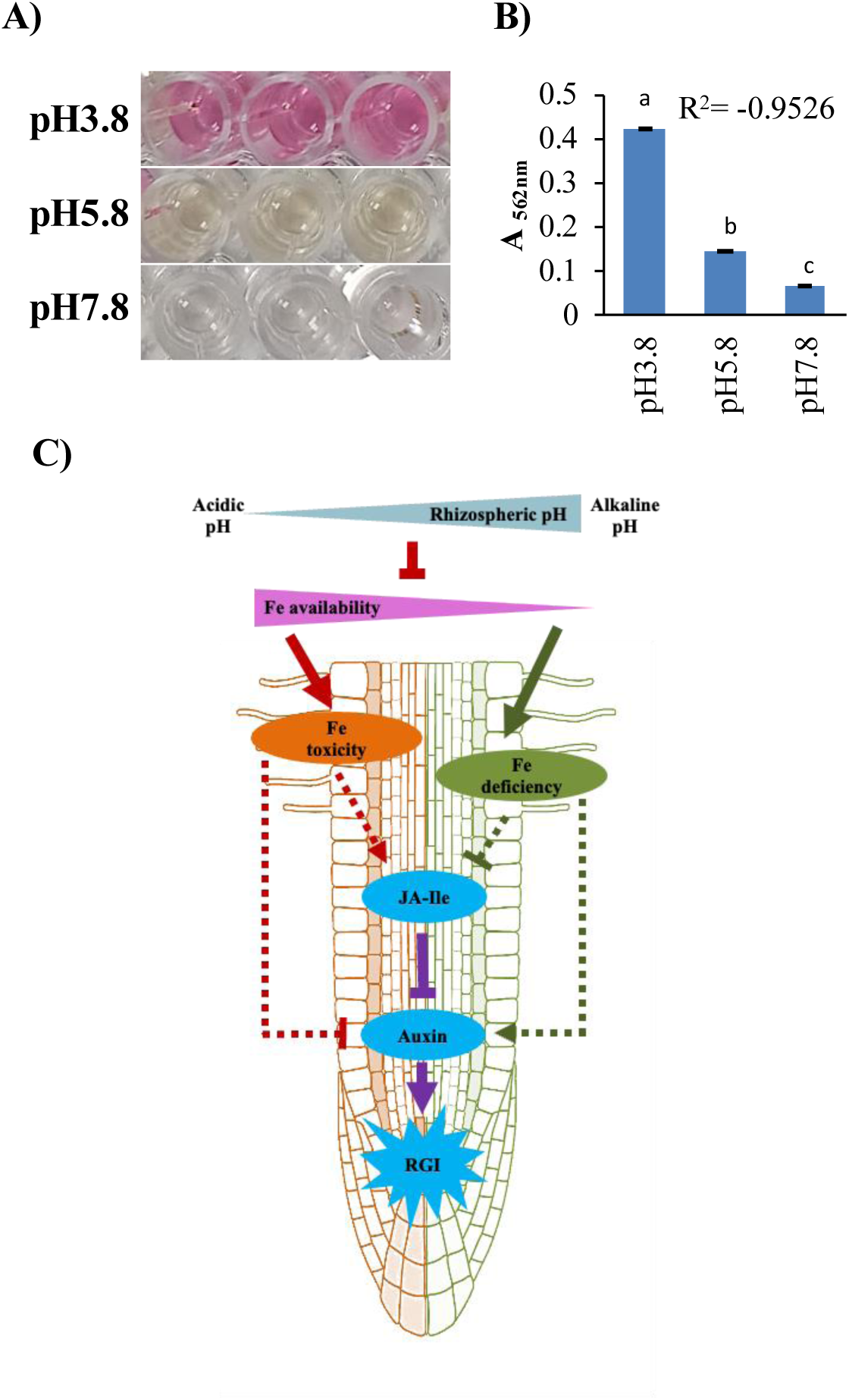
Rhizospheric pH modulates Fe availability in the media. **A)** Ferrozine assay showing the availability of Fe^2+^ (pink color) in the unutilized media over the rhizospheric pH range. **B)** Spectrophotometric quantification (absorbance at 562nm) of Fe(II)-ferrozine complex showing decreasing Fe^2+^ concentration on increasing rhizospheric pH. Each data point indicate the mean value (n=3) with SE among the replicates. R^2^ indicates the person correlation coefficient between the average absorbance (A_562nm_) and the corresponding pH values. Significance was analyzed using ANOVA, groups with different letters differ significantly based on Benjamini–Hochberg-adjusted post hoc pairwise *t*-tests (*P* < 0.05). **C)** A working model for pH mediated RGI showing increase in rhizospheric pH inversely regulate Fe bioavailability in the media creating Fe toxicity during acidic and Fe deficiency during alkaline pH conditions. The Fe toxicity during acidic pH favors induction of JA-Ile and inhibits auxin accumulation in rice roots. On the other hand, alkaline pH mediated Fe deficiency inhibits JA-Ile and promotes auxin accumulation in rice root. While JA can antagonistically regulate auxin signaling, the relative auxin accumulation in root act as the key player in regulating RGI pattern against rhizospheric pH.

## Discussion

Rhizospheric pH impacts plants fitness in several ways including nutrient imbalances, growth processes and microbiome activity in the rhizosphere (**Gregory & Hinsinger, 1999**; **Rousk et al., 2010; Fierer and Jackson, 2006**). Despite of known severe effects on plants fitness, the mechanistic framework of pH_Ext_ perception in plant is unknown (**Tsai and Schmidt, 2021**). From this context our previous study identified RGI as the primary response against severe pH_Ext_ conditions where spatial auxin abundance was demonstrated to be responsible for the RGI over the entire range of pH_Ext_ (**Singh et al., 2025**). The study also revealed that jasmonate acts as antagonistic upstream regulator of auxin. Additionally, pH is known to influences RSA by altering auxin transport and distribution, thereby affecting lateral root formation, cell expansion and branching patterns (**Barbez et al., 2017; Bibikova et al., 1998**; **Monshausen et al., 2007**; **Lager et al., 2010; Singh et al., 2025**). While most of these studies were only limited to the model plant Arabidopsis having tap-root-system, no information was available on effect of pH_Ext_ in crops like rice representing fibrous-root-system architecture. These differences made us to investigate how rice responds to changing pH_Ext_ where we observed that alike Arabidopsis, rice also showed root growth inhibition as a typical response towards both the extremes of pH_Ext_ (**Figure 1; Singh et al., 2025**). In the current study, we have also demonstrated that the physiological response against pH_Ext_ is highly conserved among Arabidopsis and rice. Similar to Arabidopsis, increasing pH_Ext_ was associated with decreasing JA accumulation (**Figure 4G**) and increasing auxin accumulation (**Figure 4H**) in rice roots. In addition to the common RGI pattern on acidic and alkaline pH, the micro-phenotypic observations like lateral root elongation (**Figure 1H-J**) on increasing pH_Ext_ also supported systemic increase in auxin abundance/signaling on increasing pH_Ext_ values (**Rahman et al., 2002**; **Wang et al., 2003**; **Nakamura et al., 2023**). Further, the rescue of acidic pH mediated RGI on auxin treatment and alkaline pH mediated RGI on MeJA treatment validated that the RGI is majorly governed by these two phytohormones (**Figure 5E-H**). The analysis with previously published transcriptome datasets on auxin and JA treatment responses in rice roots confirmed that JA signaling acts antagonistic to auxin validating the notion that Arabidopsis and rice both have similar phenotypic as well as physiological responses against pH_Ext_ (**Figure S1A-B**).

These physiological and developmental reorganizations though have strengthened the idea that plants do perceive and responds to pH_Ext_, but the mechanism of pH perception in plants is not known (**Tsai and Schmidt, 2021**). Among the common theories of pH perception in plants, one favors pH as an integrative signal that reflects composite picture of the plant’s environment including nutrient status and microbiome. Another perspective is the distributed sensing model, in which pH perception arises from the collective activity of multiple sensors encoded by the genome (**Tsai and Schmidt, 2021**). In this context, knowing that plants have dedicated and fine-tuned nutrient sensing mechanisms which can even distinguish among the isoforms of the elements, the perspective that the changes in elemental bioavailability due to change in pH_Ext_ was highly convincing. Considering influence of pH_Ext_ on nutrient availability and that the plant absorbs most of the nutrients from the rhizosphere, we analyzed if there is any change in micronutrient assimilation on different pH_Ext_ values. The root specific ICPMS analysis revealed that micronutrients including Mn, Mg, Co, Cu and Zn showed increasing assimilation on increasing pH_Ext_. Interestingly, only Fe showed a reverse pattern i.e. increased assimilation on acidic pH and reduced assimilation on alkaline pH (**Figure 5A**). In search of the mechanism for pH_Ext_ sensing in rice (if it is regulated by external stimuli), we wanted to explore the factors which can be justified on a broader pH range and can also be correlated with the plant phenotypic and physiological changes i.e. root length and transcriptomic status. Alkaline pH_Ext_ has been previously shown to regulate Fe deficiency related transcriptome in Arabidopsis (**Tsai and Schmidt, 2020).** Interestingly, **Feng et al., 2024** has very recently reported involvement of Fe toxicity as the cause of RGI on acidic pH_Ext_. Our data confirmed Fe toxicity and deficiency among the key responses against acidic and alkaline pH conditions, respectively (**Mahender et al., 2019**). While both Fe toxicity and deficiency are well known for negatively impact root growth in rice (**Fageria et al., 2008; Sun et al., 2017**), our data validated that the plants experienced higher Fe assimilation on acidic pH and Fe deficiency on alkaline pH which perfectly justified the RGI pattern on either side of the optimal pH (**Figure 5**).

Fe availability seems to be the most important factor in terms of the pH_Ext_ perception because Fe toxicity and deficiency not only justified the phenotypic observations but also physiological responses. First, the increase in auxin abundance and decrease in JA abundance almost followed a linear pattern on increasing pH_Ext_, similarly the Fe availability pattern in the media was also observed with a very good negative correlation (R^2^= -0.9526) with increasing pH values (**Figure 7A-B**). Second, the single factor is applicable to a broader range of pH which can be deduced deferentially by the plants. For example, higher Fe availability during acidic pH results into Fe toxicity response favoring JA signaling while lower Fe availability during alkaline pH results into Fe deficiency response favoring auxin signaling in the plants. Third, the two critical regulators of RGI against pH_Ext_ (i.e. JA and auxin) can be regulated differentially by Fe toxicity/deficiency responses. Also, the antagonistic crosstalk between JA and auxin intensifies the response. Finally, Fe has been emerged as the perfect external stimulus which is proportionally regulated by the pH_Ext_ and deferentially deduced by the plants into the typical physiological response in the form of JA-auxin crosstalk which can further defines the downstream phenotypic responses like RGI, lateral root elongation etc. The observations were observed to be critically related to Fe toxicity and Fe deficiency as induced auxin signaling during Fe deficiency may be an adaptive strategy for enhancing Fe uptake by promoting lateral root elongation, particularly under low-Fe conditions. Similarly, Fe toxicity mediated induction of jasmonates also fits into the perspective that induced JA signaling protects plants against Fe toxicity by inhibiting root development as well as by downregulating the genes responsible for iron uptake. This hypothesis is particularly relevant when multiple bHLH transcription factors are already known to interact with FIT to promote JA mediated inhibition of Fe uptake in Arabidopsis (**Cui et al., 2018; Maurer et al., 2011**).

The study, therefore, strongly validated that in response to changing pH conditions the differential bioavailability of Fe is decoded by the plants and that the Fe toxicity and/or Fe deficiency responses participate in activating the relevant physiological responses (**Figure 7C**). Although the RGI is regulated via Fe toxicity/deficiency response, the JA treatment mediated rescue of RGI without restoring Fe assimilation in roots, suggested the RGI response to be Fe independent. Therefore, Fe seems to be involved in pH sensing, and the RGI seems to be governed by the JA-auxin crosstalk. The future research in this direction may identify the molecular players integrating Fe toxicity/deficiency response to JA-auxin crosstalk and therefore may provide valuable leads towards generating pH tolerant/insensitive cultivars.

## Acknowledgements and funding

The research was supported by the MK Bhan Young Researcher Fellowship and Grant, DBT, India. We acknowledge NABI research facilities including microscopy and ICPMS.

## Author contributions

APS conceptualized the study and designed experiments. APS, HB, RG performed experiments. APS, AKP analyzed data and wrote the manuscript. All authors contributed to reviewing the manuscript.

## Data availability

The data supporting the findings of this study is available within the article. The transcriptomic data generated in this study have been deposited in the NCBI Sequence Read Archive (SRA) under BioProject accession number PRJNA1547916.

## Supporting Information

**Fig. S1. Heatmap demonstrating crosstalk between auxin and JA signaling.**

**Fig. S2. Rhizospheric pH regulates JA-auxin crosstalk via Fe deficiency response.**

**Fig. S3. Perl’s staining showing Fe assimilation in rice roots under Fe sufficient and Fe deficient conditions along with MeJA treatment.**

**Fig. S4. Strategy-II for Fe acquisition is functional but inefficient in rice on alkaline rhizospheric pH.**

**Table S1. List of RNA-seq datasets used in PCA analysis among rhizospheric pH and hormone treatment responses**

**Table S2. List of RNA-seq datasets used in PCA analysis among rhizospheric pH, Fe deficiency and toxicity responses.**

**Table S3. List of primers used for qRT-PCR analysis in the study.**

## References

Arnon, D. I. and C. M. Johnson. (1942) Influence of Hydrogen Ion Concentration on the Growth of Higher Plants Under Controlled Conditions. Plant Physiol. 17: 525–539.

Aung, M.S. and Masuda, H. (2020) How Does Rice Defend Against Excess Iron?: Physiological and Molecular Mechanisms. Front Plant Sci. 11:1102.

Bailey, M., Hsieh, E.J., Tsai, H.H., Ravindran, A. and Schmidt, W. (2023) Alkalinity modulates a unique suite of genes to recalibrate growth and pH homeostasis. Front Plant Sci. 14:1100701.

Barbez, E., Dünser, K., Gaidora, A., Lendl, T. and Busch, W. (2017) Auxin steers root cell expansion via apoplastic pH regulation in Arabidopsis thaliana. Proc Natl Acad Sci U S A. 114(24):E4884–E4893.

Bibikova, T.N., Jacob, T., Dahse, I. and Gilroy, S. (1998) Localized changes in apoplastic and cytoplasmic pH are associated with root hair development in Arabidopsis thaliana. Development. 125(15):2925–34.

Boukhalfa, H. and Crumbliss, A.L. (2002) Chemical aspects of siderophore mediated iron transport. Biometals 15, 325–339.

Chen, C., Chen, H., Zhang, Y., Thomas, H.R., Frank, M.H., He, Y. and Xia, R. (2020) TBtools: An Integrative Toolkit Developed for Interactive Analyses of Big Biological Data. Mol Plant. 13(8):1194–1202.

Chen, W.W., Yang, J.L., Qin, C., Jin, C.W., Mo, J.H., Ye, T. and Zheng, S.J. (2010) Nitric oxide acts downstream of auxin to trigger root ferric-chelate reductase activity in response to iron deficiency in Arabidopsis. Plant Physiol. 154(2):810–9.

Cui, Y., Chen, C.L., Cui, M., Zhou, W.J., Wu, H.L. and Ling, H.Q. (2018) Four IVa bHLH Transcription Factors Are Novel Interactors of FIT and Mediate JA Inhibition of Iron Uptake in Arabidopsis. Mol Plant. 11(9):1166–1183.

Du, H., Wu, N., Fu, J., Wang, S., Li, X., Xiao, J. and Xiong, L. (2012) A GH3 family member, OsGH3-2, modulates auxin and abscisic acid levels and differentially affects drought and cold tolerance in rice. J Exp Bot. 63(18):6467–80.

Fageria, N.K. and Zimmermann F.J.P. (1998) Influence of pH on Growth and Nutrient Uptake by Crop Species in an Oxisol. Communications in Soil Science and Plant Analysis 29: 2675–2682.

Fageria, N.K., Santos, A.B., Barbosa Filho, M.P. and Guimarães, C.M. (2008) Iron Toxicity in Lowland Rice, Journal of Plant Nutrition, 31:9,1676–1697.

Fang, X.Z., Xu, X.L., Ye, Z.Q., Liu, D., Zhao, K.L., Li, D.M., Liu, X.X. and Jin, C.W. (2024) Excessive iron deposition in root apoplast is involved in growth arrest of roots in response to low pH. J Exp Bot. 75(10):3188–3200.

Fierer, N. and Jackson, R.B. (2006) The diversity and biogeography of soil bacterial communities. Proc Natl Acad Sci U S A. 103(3):626–31.

Gregory, P. and Hinsinger, P. (1999) New approaches to studying chemical and physical changes in the rhizosphere: an overview. Plant and Soil 211, 1–9.

Jan, R., Kim, N., Asif, S., Asaf, S., Lubna, Farooq, M., Khan, Z. and Kim, K.M. (2025) Identification and evaluation of low-pH-tolerant Cheongcheong/Nagdong-double haploid rice lines via QTL analysis. BMC Plant Biol. 25(1):525.

Kraemer, S.M. (2004) Iron oxide dissolution and solubility in the presence of siderophores. Aquat. Sci. 66, 3–18.

Lager, I., Andréasson, O., Dunbar, T.L., Andreasson, E., Escobar, M.A. and Rasmusson, A.G. (2010). Changes in external pH rapidly alter plant gene expression and modulate auxin and elicitor responses. Plant Cell Environ. 33(9):1513–28.

Liu, Q., Liang, Z., Feng, D., Jiang, S., Wang, Y., Du, Z., Li, R., Hu, G., Zhang, P., Ma, Y., Lohmann, J.U. and Gu, X. (2021) Transcriptional landscape of rice roots at the single-cell resolution. Mol. Plant. 14, 384–394.

Mahender, A., Swamy, B.P.M., Anandan, A., Ali, J. (2019) Tolerance of Iron-Deficient and - Toxic Soil Conditions in Rice. Plants (Basel). 8(2):31.

Marschner, P. (2012) Marschner’s Mineral Nutrition of Higher Plants: Third Edition. London: Academic press. doi: 10.1016/C2009-0-63043-9.

Maurer, F., Müller, S. and Bauer, P. (2011) Suppression of Fe deficiency gene expression by jasmonate. Plant Physiol Biochem. 49(5): 530–6.

Maurer, F., Müller, S. and Bauer, P. (2011) Suppression of Fe deficiency gene expression by jasmonate. Plant Physiol Biochem. 49(5):530–6.

Monshausen, G.B., Bibikova, T.N., Messerli, M.A., Shi, C. and Gilroy, S. (2007) Oscillations in extracellular pH and reactive oxygen species modulate tip growth of Arabidopsis root hairs. Proc Natl Acad Sci U S A. 104(52):20996–1001.

Nakamura, A., Hirota, Y., Shigihara, M., Watanabe, M., Sato, A., Tsuji, H. and Shimada, Y. (2023) Molecular and cellular insights into auxin-regulated primary root growth: a comparative study of Arabidopsis and rice. Biosci Biotechnol Biochem. 87(10):1145–1154.

Ortas, I. and Rowell, D.L. (2004) Effect of Ammonium and Nitrate on Indigenous Mycorrhizal Infection, Rhizosphere pH Change, and Phosphorus Uptake by Sorghum. Communications in Soil Science and Plant Analysis, 35(13–14), 1923–1944.

Pandey, A., Devi, L.L., Gupta, S., Prasad, P., Agrwal, K., Asif, M.H., Pandey, A.K., Bandyopadhyay, K. and Singh, A.P. (2024) Jasmonate signaling modulates root growth by suppressing iron accumulation during ammonium stress. Plant Physiol. 196(4): 2213–2231.

Qian, D., Li, T., Zheng, C., Niu, Y., Niu, Y., Li, C., Wang, M., Yang, Y., An, L. and Xiang, Y. (2024) Actin-depolymerizing factors 8 and 11 promote root hair elongation at high pH. Plant Commun. 5(3):100787.

Rahman, A., Hosokawa, S., Oono, Y., Amakawa, T., Goto, N. and Tsurumi, S. (2002) Auxin and ethylene response interactions during Arabidopsis root hair development dissected by auxin influx modulators. Plant Physiol. 130(4):1908–17.

Rousk, J., Bååth, E., Brookes, P.C., Lauber, C.L., Lozupone, C., Caporaso, J.G., Knight, R. and Fierer, N. (2010) Soil Bacterial and Fungal Communities across a pH Gradient in an Arable Soil. ISME J. 4: 1340–1351.

Schenkeveld, W.D., Schindlegger, Y., Oburger, E., Puschenreiter, M., Hann, S. and Kraemer, S.M. (2014) Geochemical processes constraining iron uptake in Strategy II Fe acquisition. Environ Sci Technol. 48(21):12662–70.

Singh, A.P., Kanwar, R. and Pandey, A.K. (2025) Jasmonates Regulate Auxin-Mediated Root Growth Inhibition in Response to Rhizospheric pH in Arabidopsis thaliana. Plant Cell Environ. 48(1):756–767.

Singh, A.P., Mani, B. and Giri, J. (2021) OsJAZ9 is involved in water-deficit stress tolerance by regulating leaf width and stomatal density in rice. Plant Physiol Biochem. 162:161–170.

Singh, A.P., Pandey, B.K., Deveshwar, P., Narnoliya, L., Parida, S.K. and Giri, J. (2015) JAZ Repressors: Potential Involvement in Nutrients Deficiency Response in Rice and Chickpea. Front Plant Sci. 6:975.

Singh, A.P., Pandey, B.K., Mehra, P., Heitz, T. and Giri, J. (2020) OsJAZ9 overexpression modulates jasmonic acid biosynthesis and potassium deficiency responses in rice. Plant Mol Biol. 104(4-5):397–410.

Sun, H., Feng, F., Liu, J. and Zhao, Q. (2017) The Interaction between Auxin and Nitric Oxide Regulates Root Growth in Response to Iron Deficiency in Rice. Front Plant Sci. 8:2169.

Tian, H., Xu, L., Li, X. and Zhang, Y. (2025) Salicylic acid: The roles in plant immunity and crosstalk with other hormones. J Integr Plant Biol. 67(3):773–785.

Tsai, H.H. and Schmidt, W. (2020) pH-dependent transcriptional profile changes in iron-deficient Arabidopsis roots. BMC Genomics. 21(1):694.

Tsai, H.H. and Schmidt, W. (2021) The enigma of environmental pH sensing in plants. Nat Plants. 7(2):106–115.

Ueda, Y., Yamaji, N. and Wissuwa, M. (2025) Comparative Transcriptome Reveals ART1-Dependent Regulatory Pathways for Fe Toxicity Response in Rice Roots. Physiol Plant. 177(4):e70398.

von Uexküll, H.R. and Mutert, E. (1995) Global extent, development and economic impact of acid soils. Plant Soil 171, 1–15.

Von Wiren, N., Marschner, H. and Romheld, V. (1996) Roots of Iron-Efficient Maize also Absorb Phytosiderophore-Chelated Zinc. Plant Physiol. 111(4):1119–1125.

Wairich, A., de Oliveira, B.H.N., Arend, E.B., Duarte, G.L., Ponte, L.R., Sperotto, R.A., Ricachenevsky, F.K. and Fett, J.P. (2019) The Combined Strategy for iron uptake is not exclusive to domesticated rice (Oryza sativa). Sci Rep. 9(1):16144.

Wang, S., Taketa, S., Ichii, M., Xu, L., Xia, K. and Zhou, X. (2003) Lateral root formation in rice (*Oryza Sativa* L.): differential effects of indole-3-acetic acid and indole-3-butyric acid. Plant Growth Regulation 41, 41–47.

Wasternack, C. and Hause, B. (2013) Jasmonates: biosynthesis, perception, signal transduction and action in plant stress response, growth and development. An update to the 2007 review in Annals of Botany. Ann Bot. 111(6):1021–58.

Yang, G., Metze, H.S. and Krämer, U. (2026) Cross-Species Plasticity and Divergence Under Alkaline Soil Signature Conditions Differentiating PYE From FIT Target Transcripts in the Arabidopsis thaliana Ferrome. Plant Cell Environ. 49(9):6719–6736.

Yi, Y. and Guerinot, M.L. (1996) Genetic evidence that induction of root Fe(III) chelate reductase activity is necessary for iron uptake under iron deficiency. Plant J. 10(5):835–44.

Yu, Y., Zhang, H., Long, Y., Shu, Y. and Zhai, J. (2022) Plant Public RNA-seq Database: a comprehensive online database for expression analysis of ∼45 000 plant public RNA-Seq libraries. Plant Biotechnol J. 20(5):806–808.

Zhang, H., Yu, F., Xie, P., Sun, S., Qiao, X., Tang, S., Chen, C., Yang, S., Mei, C., Yang, D., Wu, Y., Xia, R., Li, X., Lu, J., Liu, Y., Xie, X., Ma, D., Xu, X., Liang, Z., Feng, Z., Huang, X., Yu, H., Liu, G., Wang, Y., Li, J., Zhang, Q., Chen, C., Ouyang, Y. and Xie, Q. (2023) A Gγ protein regulates alkaline sensitivity in crops. Science 379(6638):eade8416.

