## Supplemental Figures for "Iron sensing modules participate in rhizospheric pH perception in rice"

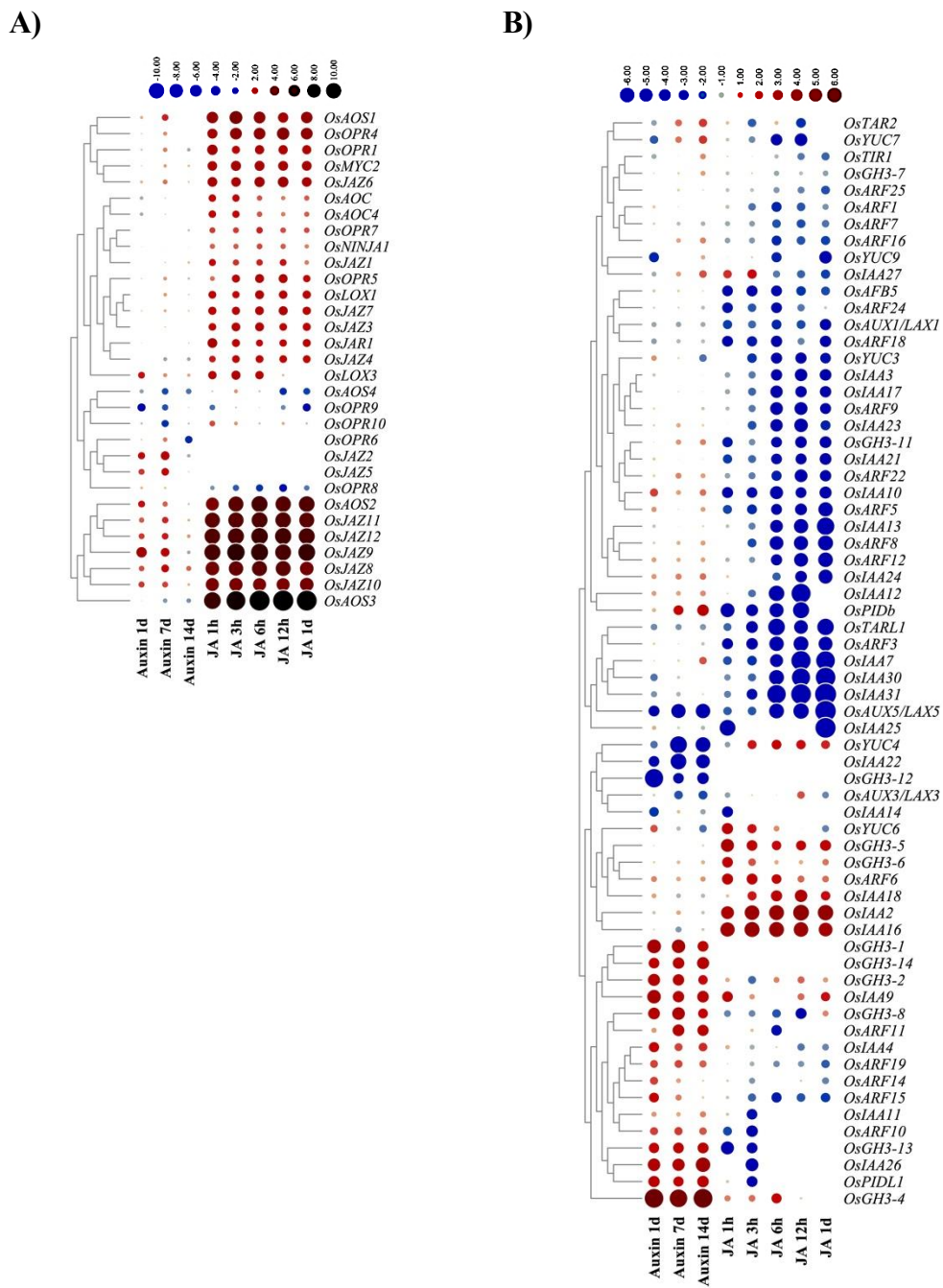

**Figure S1. Heatmap demonstrating crosstalk between auxin and JA signaling. A)** Expression pattern of jasmonate signaling/biosynthesis genes on auxin treatment and JA treatment among different RNA-seq datasets. **B)** Expression pattern of auxin signaling/biosynthesis genes on auxin treatment and JA treatment in the same datasets. Genes with expression $\text{Log}_2\text{FC} > 0.5$ in at least one of the RNA-seq data was considered for plotting.

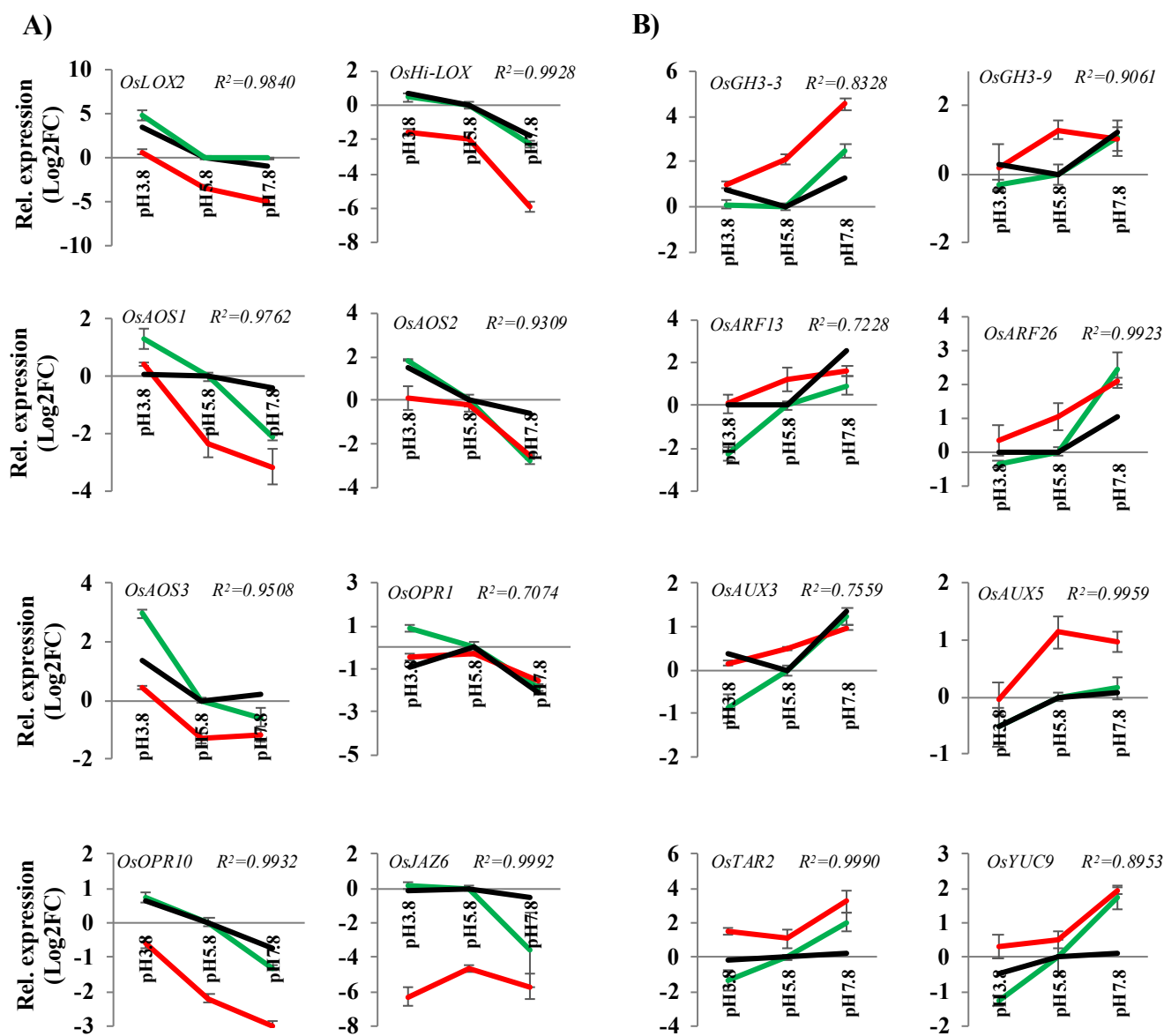

**Figure S2. Rhizospheric pH regulates JA-auxin crosstalk via Fe deficiency response.** qRT-PCR analysis showing effect of rhizospheric pH on **A)** JA and **B)** auxin signaling/biosynthetic genes expression in rice roots under Fe sufficient (green) and Fe deficient conditions (red). The black lines indicates the expression of the respective genes observed in the transcriptome data (Fe sufficient conditions) over different pH conditions. Each datapoint represents mean of three biological replicates with error bars representing SE among the replicates. $R^2$ value shows the Pearson correlation coefficient among the mean of qRT-PCR data (Fe sufficient condition) and transcriptome data over different pH conditions.

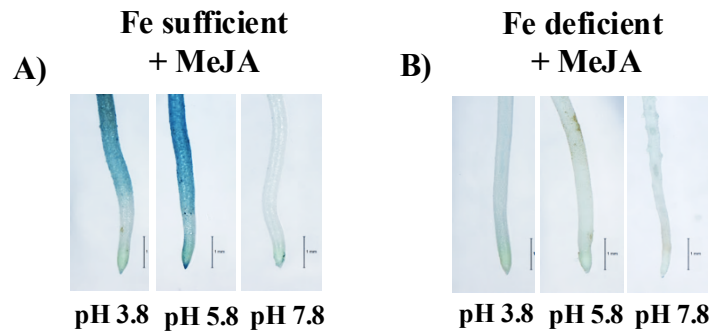

**Figure S3: Perl's staining showing Fe assimilation in rice roots under A) Fe sufficient and B) Fe deficient conditions with different rhizospheric pH and MeJA (0.01μM) treatment, simultaneously.**

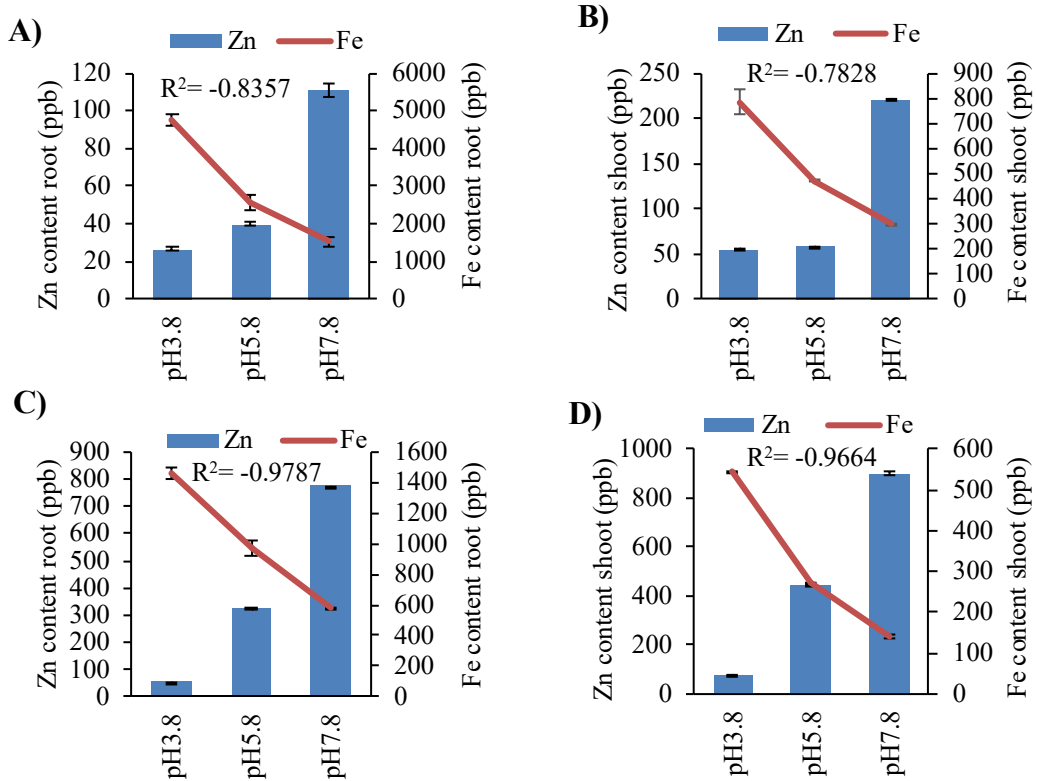

**Figure S4. Strategy-II for Fe acquisition is functional but inefficient in rice on alkaline rhizospheric pH.** Effect of rhizospheric pH on Zn and Fe assimilation in root (A) and shoot (B) under Fe sufficient conditions and root (C) and shoot (D) under Fe deficient conditions. Each datapoint represent mean of three independent replicates with error bars represents SE among the replicates. Pearson correlation coefficient ($R^2$) was calculated between Iron (Fe) and Zinc (Zn) assimilation in respective tissues during differential rhizospheric pH conditions.
